# Light dependent morphological changes can tune light absorption in iridescent plant chloroplasts

**DOI:** 10.1101/2020.09.01.277616

**Authors:** Miguel A. Castillo, William P. Wardley, Martin Lopez-Garcia

**Affiliations:** NATURAL AND ARTIFICIAL PHOTONIC STRUCTURES AND DEVICES GROUP, INTERNATIONAL IBERIAN NANOTECHNOLOGY LABORATORY-INL, BRAGA, PORTUGAL

## Abstract

Chloroplasts, the organelles responsible for photosynthesis in most plants and algae, exhibit a variety of morphological adaption strategies to changing light environments which can have important yet overlooked light scattering effects. This can be even more significant for iridoplasts, specialized chloroplasts whose tissue is arranged as a photonic multilayer producing a characteristic strong blue reflectance associated to a wavelength selective absorption enhancement relevant for photosynthesis.

In this work, we study how the photonic properties of iridoplasts are affected by light induced dynamic changes using realistic data extracted from previous reports. Our results show a reflectance red-shift from blue to green under increasing light intensity. Consequently, the light absorption enhancement induced by the photonic nanostructure is also redshifted. We also show that the photonic properties are resilient to biologically realistic levels of disorder in the structure. We extended this analysis to another photonic nanostructure-containing chloroplast, known as a bisonoplast, and found similar results, pointing towards similar properties in different plant species. We finally found that all types of chloroplasts can tune light absorption depending on light conditions. In general, our study opens the door to understanding how dynamic morphologies in chloroplasts can affect light scattering and absorption.

## 1. Introduction

The vast majority of life on Earth, and certainly all higher plant and animal species, ultimately derive all their energy from photosynthesis; the conversion of solar energy, carbon dioxide and water into carbohydrates. In high plants, algae and cyanobacteria, this process is carried out in chloroplasts. Due to their crucial role in energy harvesting, their composition and morphology have been investigated to a great extent. The arrangements of protein and pigment constituents in the thylakoid membranes, such as Photosystems I and II, and, to a large extent, the molecular processes by which photons are absorbed by chloroplast pigments and transmitted to the reaction centres are well understood. This has led to very relevant discoveries such as the extreme exciton transport quantum efficiency in PS complexes (1,2). However, little attention has been paid to the role of the tissue’s structural arrangement and consequent light scattering effects within chloroplasts. A recent theoretical work (3) has suggested that the stratified morphology of thylakoids within the chloroplast, known as grana (see Figure 1), could enhance light absorption through interference. Interestingly, the novel concept that thylakoid arrangement could influence light absorption in PS through nanophotonic-like arrangements had been reported previously for a specific type of specialized chloroplast, known as an iridoplast (4).

**Figure 1.**
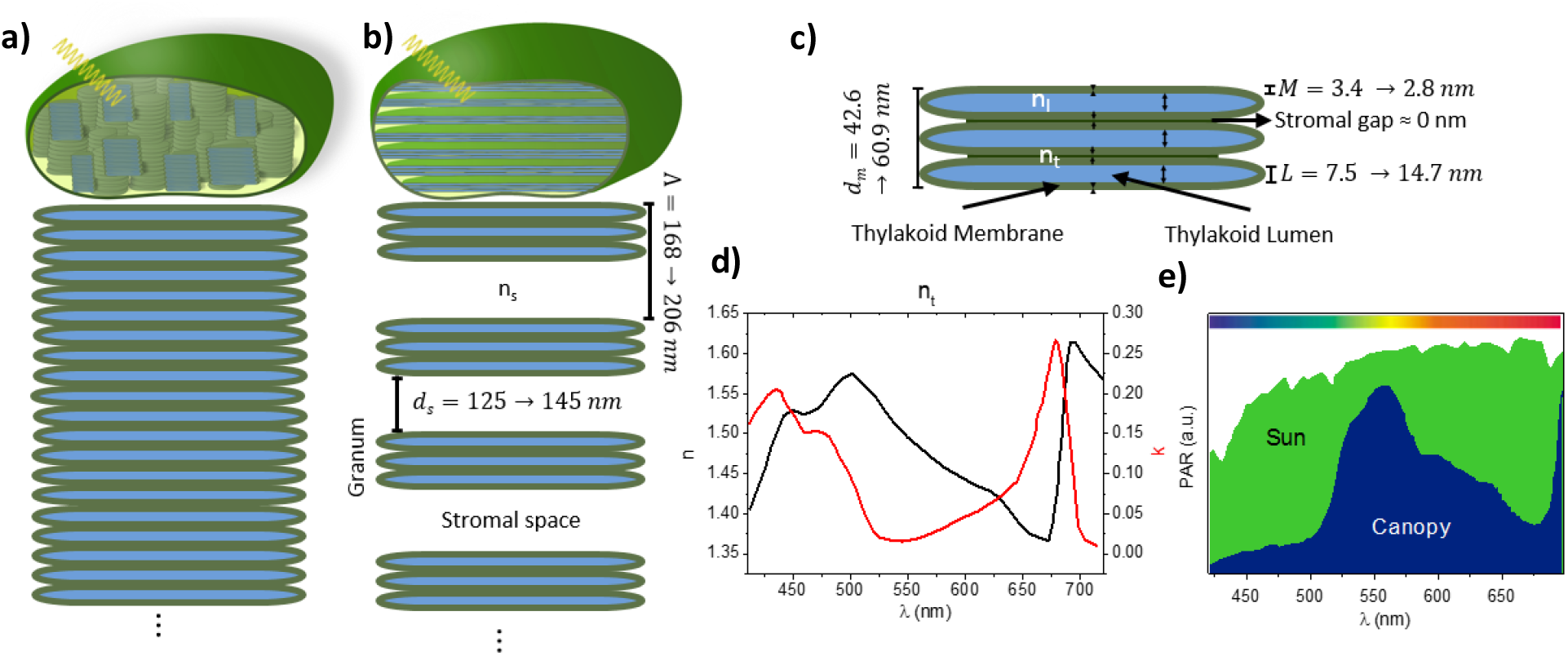
Photonic structure of the chloroplast and iridoplast. **a)** Schematic of a chloroplast showing the distribution of the layers. **b)** Grana distribution in an iridoplast. Note the expansion if the stroma and grana going from low to high light conditions. **c)** Detailed granum composition and layer thicknesses under low and high light conditions. **d)** Refractive index of the thylakoid membrane (27). **e)** Normalized photosynthetically active radiation (PAR) spectra under direct sun (green; i.e. high light) and forest canopy shade (blue; i.e. lowlight) (5). We note that these two spectra are not to scale to each other.

Chloroplasts in plants and algae can experience highly variable light environments, affecting both the quality and quantity of light, such as the available wavelengths and their intensity (5). Therefore, to avoid photodamage, chloroplasts hold several molecular mechanisms to adjust their response to a changing light environment (6). Moreover, chloroplasts also undergo morphological changes during adaptation to different light environments (7). For example, electron microscope studies have shown that, upon illumination, dark-adapted grana will contract, flatten and become more ordered (8,9). This remodelling has been linked to photodamage protection in the past (10,11). More recent studies looked at the nanostructural changes of thylakoids under increasing light intensity (12) and found that the grana volume increases under high light, contradicting previous findings. Taking into account the advances in electron microscopy treatments to biological tissue, we believe that the more recent studies are more reliable regarding spatial resolution.

Light induced structural changes of chloroplasts have been the target of much research (13–16), nevertheless their mechanism and biological functions remain unknown. Recent research suggests that the shift to an expanded (contracted) system at high (low) light conditions could help (inhibit) protein diffusion (12) and cyclic electron transfer (17), creating a photodamage protection mechanism. It also inhibits (helps) linear electron transfer, improving photosynthetic efficiency (15,17). This regulatory system is known as a state transition (18). Dark adaptation induces reduction of the lumen thickness, allowing for inter-thylakoid chemical exchange (12). To date however, very few studies are available regarding the morphological adaptations of thylakoid membranes in the chloroplast. Interestingly, recent modelling results suggested that the grana expansions could also increase light absorption (3). However, this model is probably oversimplified since the layers were all uniformly expanded which does not agree with the experimental evidence (12).

Iridoplasts are characteristic of several species of the genus *Begonia*, a set of shade dwelling plants whose leaves show strong blue coloration under low light conditions (4). They are specialised epidermal cell chloroplasts presenting a characteristic structural arrangement which provides them a strong blue iridescence. Contrary to non-iridescent chloroplasts, iridoplasts present a persistent number of thylakoids in each granum. Simultaneously, each grana is separated from the next by a uniform distance, forming a Bragg-stack like structure with photonic properties (19). Preliminary works by Gould et al. (20) suggested that the structural colour of the iridoplast was a cue for pollinators or to deter predators. However, the most recent advances have shown that the most likely function of the photonic structure of iridoplasts is to selectively enhance light absorption in heavily attenuated light environments (where these species are usually found) and even enhance the quantum yield at the thylakoid membrane (4). However, in order to fully understand the effects of these specialized photonic structures and their biological functions for the plant, it is important to consider the highly variable light environments to which plants usually have to adapt and the mechanisms that chloroplasts and iridoplasts could make use of to adapt the photonic response to their environment.

In several *Begonia sp*. that grow iridoplasts, the development of highly ordered thylakoid arrangements with intrinsic photonic properties in iridoplasts is highly plastic during the early stages of leaf growth. For a specimen grown under high light conditions, iridoplasts present morphologies similar to the thylakoid arrangement in a non-photonic chloroplast while specimens grown under low light or deep-shade light conditions, the iridoplast develops thylakoid and granum arrangements with photonic crystal properties (21,22). Herein, we focus on the latter case where the plant has been grown under low light conditions. Since short- and long-term light sensitive structural changes can occur after leaf development (16,23,24) we focus at how short time changes (timeframe < 2 min in the natural system (12)) might affect the photonic properties of iridoplasts whilst developmental photonic properties will be studied elsewhere.

Bisonoplasts represent another specialised chloroplast. This organelle is also found on deep shade dwelling plants, such as the *Selaginella erythropus* (26). The arrangement of the thylakoid membranes in bisonoplasts consist of a well-defined two phase structure, which gives it its name. Away from the microphyll surface, the lower phase presents a structure similar to the standard non photonic chloroplast distribution. Near the microphyll surface, the upwards phase presents a highly ordered structure similar to the iridoplast. Outside of shade-dwelling plants, other photonic structures that play a role in photosynthesis are also found in some species of giant clams. They have biophotonic structures with multilayer structures which also act as Bragg mirrors and have been found to reduce photodamage by spatially redistributing the available sunlight (34).

In this work, we use iridoplasts as a first step to understand the functions of the morphological adaptations over light absorption in chloroplasts. Since iridoplasts present characteristic photonic properties that are strongly dependent on the morphology, they represent an ideal case study. Moreover, iridoplasts are known to present dynamic mechanisms of adaptation to the light environment. The leaves of the plant species *Begonia pavonina* reversibly change between blue and green coloration under low and high light conditions respectively. However, the biological advantage and the mechanism remain unknown (4). In this paper, we assume that documented light sensitive morphological changes in chloroplasts can be extrapolated to iridoplasts. We use numerical methods to quantify the effects over structural colour and light absorption of iridoplasts when they undergo similar changes such as those observed for chloroplasts (12). We also take structural changes from an iridoplast from a different *Begonia* species (25) and assume these also occur in blue iridoplasts of the *Begonia pavonina*. We start by calculating how these changes affect the angle dependent reflection of iridoplasts and the absorption compared to non-structured chloroplasts. We then repeat these for a more biologically realistic case, where disorder in the thicknesses is considered. We extend this technique and perform a similar analysis for the bisonoplast. We finish by comparing how light sensitive structural changes affect absorption of photonic and non-photonic chloroplasts individually throughout the visible and near infra-red spectral range.

## 2. Methods

From a photonics perspective, chloroplasts can be modelled as a multilayer stack (see Figure 1 a) formed by the repetition of a unit cell (granum) consisting of three distinct tissues: thylakoid lumen (*L*), thylakoid membrane (*M*) and stromal gap (Figure 1 a, c). Each granum is composed of a variable number of thylakoids (*N*_*t*_) depending on whether they were grown under low and high light conditions (17). It is well established that the total number of grana per chloroplast (*N*_*g*_) increases from low light to high light making *N*_*g*_ a highly variable parameter.

Iridoplasts (Figure 1 b) present a regular number of thylakoids per grana (2 ≤ *N*_*t*_ ≤ 4) (4). Contrary to chloroplasts though, iridoplasts characterize themselves by presenting granum regularly separated from each other by a well-defined stromal space of length *d*_*s*_. Compared to chloroplasts, the stromal space between each grana is considerably larger with *d*_*s*_ > 100 nm (4). We consider that both iridoplasts and chloroplasts consist of seven unit cells (*N*_*g*_ = 7) plus one stack of granum so that the model begins and ends with the granum. Both the incident media and the medium at which the transmission is calculated are stroma. Note that in chloroplasts a stromal gap is known to also exist between thylakoids of the same grana but this distance is usually very small (around 3.6 nm (12)), making its characterization under TEM very challenging or even impossible. This gap has not been identified experimentally for iridoplasts either, most likely due to the lack of resolution of the cryo-TEM tools used for their analysis (4). Since numerical results show the stromal gap has only a small impact in the photonic properties of the system (see Figure A 1) for values >3 nm, we will neglect it in this work. Finally *d*_*s*_ is preserved in the whole length of the iridoplasts as shown in the sketch of Figure 1 b. Given that there is also a regular number of grana per iridoplast (6 < *N*_*g*_ < 8), we can model a chloroplast as a particular case of iridoplast with *d*_*s*_ = 0 and *N*_*g*_^*chlo*^ = *N*_*g*_^*irid*^. We will use this definition several times along this work to make the photonic response of iridoplast and chloroplast comparable.

Different plant species present slightly different structures of chloroplasts, therefore one could expect their light sensitive behaviour to be different as well. This is clear in ref. (25) where the stromal gap from iridoplasts of different *Begonia* species present very distinct reactions to light conditions. Nonetheless, in our optical model we assume that the layers of the iridoplast considered here will undergo the same proportional dynamic changes as non-structured chloroplasts in ref. (12) and blue iridoplasts in ref. (25), where in the latter the stroma of the blue iridoplast of *Begonia rockii* has been measured to expand (25) when going from low to high light conditions.

Building upon previous studies (4,12,25), we have summarized in Table 1 the expected thicknesses under high and low light for the different membranes forming the iridoplast. Table 1 shows the values for the ideal thicknesses (*μ*) and light sensitive variations with their respective uncertainties (standard deviation: *σ*), which are crucial to model more realistic scenarios where disorder occurs. In this paper, we will first introduce a perfect scenario with no thickness uncertainty (*σ = 0*) followed by a more realistic case where a given amount of thickness variability is considered (*σ ≠ 0*).

**Table 1:** Average thickness (*μ*) and uncertainties (*σ* the different photosynthetic layers of iridoplasts un different light conditions and the variation of each la when going from dark to high light.

|  | Thickness under low light (nm) (4) | Variation going to high light (%) (12,25) | Thickness under high light (nm) |
| --- | --- | --- | --- |
| Thylakoid Membrane (M) | $3.4 \pm 0.8$ | $83 \pm 6$ | $2.8 \pm 0.7$ |
| Thylakoid Lumen (L) | $7.5 \pm 0.8$ | $196 \pm 40$ | $14.7 \pm 3.4$ |
| Stromal gap ($d_s$) | $125 \pm 16$ | $116 \pm 5$ | $145 \pm 19.6$ |

The optical properties of chloroplasts and iridoplasts are calculated by the Transfer Matrix Method (TMM) using parameters in Table 1. The calculations were performed considering that lumen and stroma are aqueous media with both having the same refractive index as water (*n*_*l*_ = *n*_*s*_ = 1.33) and assumed to be constant for all wavelengths (no material dispersion). The refractive index of thylakoid membranes (*n*_*t*_) was extracted from previous studies (27). As can be seen in Figure 1 d, the refractive index presents a very strong material dispersion due to the complex pigment composition of the photosynthetic tissue. The pigment composition of iridoplasts has been largely debated but to date it has not been possible to extract them. Hence, we will assume that the optical properties of iridoplast’s thylakoid membranes are similar to those tabulated for chloroplasts. Taking into account the refractive index and morphological parameters (Table 1) of iridoplasts and chloroplasts, we can calculate the photonic properties, namely reflectance and transmission, of the multi-layer under different light conditions. Note that the refractive indices are assumed to be constant under different morphological conditions.

The absorption of iridoplasts and chloroplasts has been calculated by taking into account that *A*(*λ*) = 1 − *R*(*λ*) − *T*(*λ*), where *R*(*λ*) and *T*(*λ*) are respectively the wavelength dependent reflection and transmission of the multistack under consideration. Finally, we define the dimensionless absorptance enhancement parameter *γ*(*λ*) = *A*(*λ*)_*iridoplast*_/*A*(*λ*)_*chloroplast*_, where *A*(*λ*)_*iridoplast*_ is the absorptance of an iridoplast calculated for a given set of parameters and *A*(*λ*)_*chloroplast*_ corresponds to the absorption of the same iridoplast when *d*_*s*_ = 0. This is a powerful comparison since it allows a direct evaluation on the effects that a given structural distribution of grana can have on the overall absorption of the organelle whilst material properties are kept constant. This approach unveils those non-obvious effects of iridoplasts induced solely by the photonic properties of the structure.

The Photosynthetic Active Radiation (PAR) represents the part of the solar light spectrum available for photosynthesis in the natural environment. Thinking about the ecosystems where plant species with iridoplasts grow, we consider that low light conditions corresponds to PAR under the canopy and high light conditions show PAR under direct sun, both represented in Figure 1 e. Having found *γ*, one can calculate the total absorptance enhancement parameter (*γ*_*total*_) over the PAR under or above the forest canopy (i.e. under low or high light respectively). This is defined as a ratio between the energy absorbed with the enhancement (iridoplast) and the energy absorbed without enhancement (chloroplast, i.e. same iridoplast with *d*_*s*_ = 0):

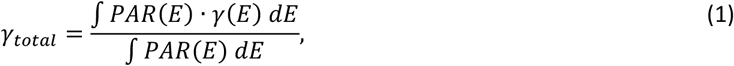

Note that both PAR and *γ* are both either at low or high light conditions. Therefore, *γ*_*total*_ > 1 corresponds to an overall light absorption enhancement and *γ*_*total*_ < 1 describes an overall light absorption reduction compared to a chloroplast. We integrate over energy rather than wavelength as the former is the biological relevant quantity for photosynthesis. With this, we investigate how iridoplasts and chloroplasts could adapt to sudden light variations.

References used to build Table 1 state that the experimental results were measured under high light with an intensity ≳ 500 μmol photons m^−2^·s^−1^ (12,21) whilst low light conditions were achieved with intensities between 15-30 μmol photons m^−2^·s^−1^ (22). Interestingly, the lowest intensities applied in the laboratory are higher than those found under the forest canopy (≃16 μmol photons m^−2^·s^−1^(5)), where those plant species showing iridoplast organelles are usually found. Also, the high intensities used in laboratory experiments are smaller than the photon count over the forest canopy at natural conditions > 700 μmol photons m^−2^·s^−1^(5)). It is then safe to assume that, by using the values in Table 1, we are analysing biologically relevant light environments.

Finally, to provide an analysis as close as possible to the natural environment, we have considered unpolarised illumination in our simulations by adding the results for P and S polarized incident light incoherently.

## 3. Results

### a. Light sensitive photonic adaptations of iridoplasts

The colour of the iridoplast, and therefore their photonic properties, have been documented to reversibly vary from blue to green under low and high light conditions respectively. However, the underlying mechanisms remain unknown (4). As a first step to model the optical properties of iridoplasts under such morphological changes, we consider the simplest case in which similar layers present the same thickness, i.e. *σ* = 0 (see Table 1). In that case, our model predicts a strong blue reflection under low light condition and normal incidence (Figure 2 a) as expected from previous reports (4). Interestingly, when parameters corresponding to high light conditions (Table 1) are applied, a strong redshift along with a strong reduction of the absolute reflectance are observed. From the colour bar in Figure 2 a, the ≈ 100 nm redshift (from 450 to 550 nm) of the dominant reflection peak will produce a switch of the structural colour of the iridoplast from blue to pale green. The redshift is a consequence of the increase in the period of the photonic crystal induced by the high light conditions. The reduction on the absolute reflection on the other hand is a consequence of the fact that the morphology under high light conditions will shift the spectrum into the lower refractive index spectra range for the thylakoid membranes (500 < *λ* < 600 nm, Figure 1d). With the same number of layers a lower refractive index will produce a weaker reflectance in that range.

**Figure 2.**
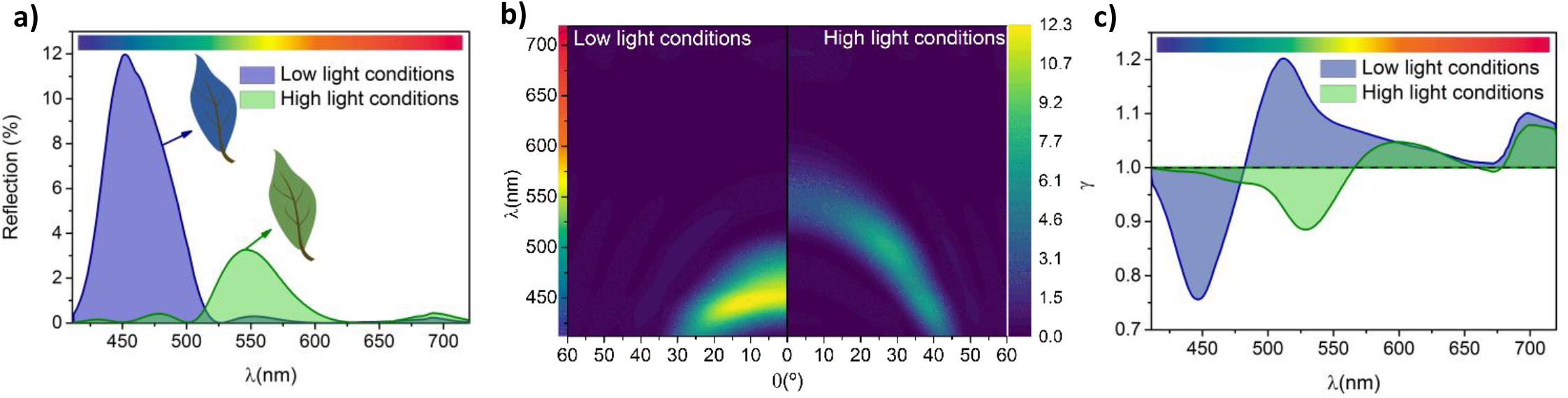
Optical properties for ordered iridoplast under changes caused by high and low light conditions. **a)** Normal incidence reflection spectrum. **b)** Angle dependent reflection spectrum. **c)** Absorption enhancement (*γ*) of iridoplasts compared to chloroplasts.

To better understand this effect, we model the evolution of the reflectance with angle (note that we are considering unpolarised incident light). Figure 3 b shows the angle (*ϑ*) dependent reflection calculated for both low and high light conditions. A clear angle dependent response is observed in both cases with a pronounced blue shift for large incident angles. This is a well-known property of structural colour produced by ordered photonic structures (28). The angular dependency of structural colour has been identified to play important roles in different biological systems, as for example the case of ref. (29) where the directional dependence of the structural coloration was found to function as a cue for pollinators. Iridoplasts and chloroplasts present a material composition with very strong dispersion (Figure 1 d). As a consequence, the maximum reflectance for low light conditions is obtained at near normal incidence. However, the reflectance under high light conditions is observed to grow at larger angles peaking at about θ ≈ 25° as a consequence of the largest *n*_*t*_ for *λ* < 500 nm. Absolute reflectance never reaches the same values as under low light conditions, which is most likely related with the non-degenerate reflectance for both polarisations under high incident angles. The bandgap of a 1D photonic crystal, such as the iridoplasts, is degenerate for both polarizations at normal incidence. That is not the case for large incident angle (i.e. high momentum) conditions (28). P-polarization will, in general, present lower absolute reflectance values and given that in these study we are considering unpolarised light, a lower absolute reflectance would be expected under high illumination. In general, Figure 2 b suggests that the characteristic strong blue reflectance of the iridoplast will shift to green under observation by the naked eye which could also in part explain why leaves of *Begonnia sp*. exposed to high light conditions do not show the characteristic blue colour of the specimens grown under low light.

**Figure 3.**
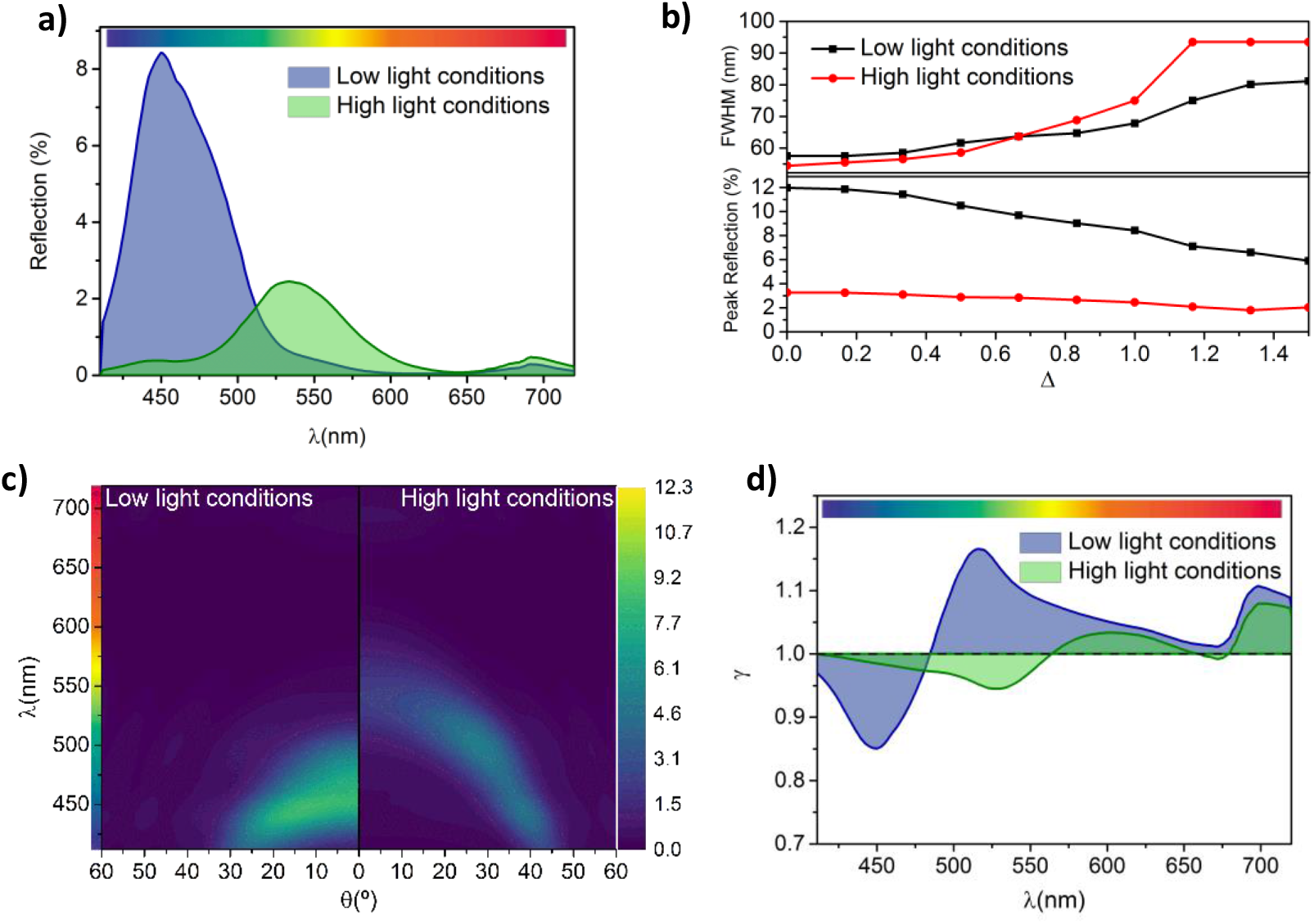
Optical properties for disordered iridoplast under dynamic changes caused by different light conditions. **a)** Normal incidence and **b)** angle dependent reflection spectrum of the iridoplast at *Δ* = 1 at low and high light conditions. **c)** Full width at half maximum (*FWHM*) and maximum reflection of the iridoplast at normal incidence for different disorder levels (i.e. varying *Δ*). **d)** Absorption enhancement (*γ*) of the iridoplast compared to a standard chloroplast both at *Δ* = 1 and at low or high light conditions.

Beyond colour production, the structure of photonic crystals can strongly influence light absorption of the iridoplast depending on the spatial position of absorbing layers. To investigate this effects we have calculated the dimensionless absorptance enhancement parameter *γ(λ)* for iridoplasts under low and high light conditions (Figure 2 c). As can be observed, *γ* presents a maximum at *λ* ≈ 510nm under low light illumination with *γ* > 1 (absorption enhancement) for *λ* ≳ 480nm. Interestingly, our model suggests that under high light conditions the expansion of the photonic crystal will flatten this parameter and produce a redshift of the γ peak to *λ* ≈ 600 nm. There also exists a redshift on the overall increase (*γ* > 1) of the absorption enhancement to *λ* ≳ 565 nm under high light conditions.

The wavelength-selective enhancement of the absorption in iridoplasts was suggested in previous reports to be related to the so called slow light effect (4). Our results in Figure 2 c show that the photonic crystal structure of the iridoplast can produce slow light effects for both high and low illumination conditions. In general, the group velocity of the light propagating is reduced near both photonic band edges (30), that is at both edges of the reflection peaks, which are at 440 nm and 520 nm in the case of low light condition. Within the band edge spectral ranges, light–matter interaction is strongly enhanced which leads to a strong absorption provided that the electric field is predominantly concentrated at the position of the absorption layers. However, given that the absorption layers in the case of iridoplasts are restricted to the thylakoid membranes and therefore to the grana, the enhancement of the absorption will only take place for the longer wavelength photonic band edge (*γ* >1).

Using equation (1), we find that *γ*_*total*_ = 1.06 under low light conditions. This means that iridoplasts absorb 6% more light solely due to its structural effects, which could be a biological significant advantage in an environment with limited light for photosynthesis. Interestingly, under high light conditions *γ*_*total*_ = 0.98 meaning the structure is not providing any advantage in terms of light absorption. In fact, for *γ*_*total*_ < 1 it is expected that chloroplasts are more effective at absorbing light than iridoplasts, therefore making the photonic crystal structure counterproductive regarding light absorption. This result could also point towards a mechanism to avoid photodamage by reducing absorption under high light conditions. Overall, our results suggest that the morphological changes that iridoplasts undergo under different light environments might have strong effects over their photonic properties and therefore over the biological functions of these organelles.

### b. Realistic Scenario

Whilst previous section have shown intriguing properties for iridoplasts under different light conditions, Table 1 shows that realistic models should include disorder in thickness. To achieve a more biologically representative scenario, we introduce disorder to the model used in the previous section.

To introduce variable morphologic inhomogeneity within our model, we introduce the degree of disorder (*Δ*). Each individual layer in the model will present a thickness (*N*_*α*_, with *α* = 1, 2… representing the thylakoid membrane, the lumen or the stromal spacing) randomly adjusted to a degree of disorder (*Δ*). We represent this as *N*_*α*_ = *μ*_*α*_ ± *δ*_*α*_, where *μ*_*α*_ represents the averaged experimentally measured thickness and *δ*_*α*_(*Δ*, σ_*α*_) represents a random number distributed normally around −*Δ* ∙ σ_*α*_ and +*Δ* ∙ σ_*α*_. Hence, *Δ* = 0 corresponds to the ideal scenario presented in the previous section and *Δ* = 1 corresponds to the expected uncertainty from reported experimental measurements. We use this for both light conditions and photosynthetic systems. The statistical dependency of the model means we have to average over a given number of simulations (*S*) to obtain the final expected reflectance. Here *S* = 50. The position of the edge of the *m*th layer is given by

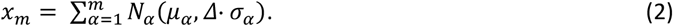

Because their positions are dependent on previous layers we can identify this variability as correlated disorder (31).

Figure 3 a shows that, compared to the ordered case (i.e. *Δ* = 0), the reflectance spectra under both illumination conditions is not significantly affected by the expected uncertainty (i.e. *Δ* =1) of the experimental data in Table 1, changing only the absolute reflectance. To evaluate the consequences of disorder in detail, we have plotted absolute values and full width at half maximum (*FWHM*) of the reflectance peaks as a function of the uncertainty parameter *Δ* (Figure 3 b). As can be observed, even though the central wavelength of the reflection peak remains mostly unchanged for both light conditions (see Figure A 2), their intensity is reduced as *Δ* increases. Increasing disorder also results in an increase of the *FWHM* which has been described in detail for artificial disorder 1D photonic crystals in the past (31). In particular, at normal incidence and under low light, the *FWHM* might vary between 55 and 80 nm and the reflection reduces from 12 to 6% for *Δ* = 0 and 1.5 respectively. Such broadening will have important implications in both the colour displayed by the iridoplast (hence the leaf) and in the properties derived from the light matter interaction enhancement due to the photonic environment in the iridoplast. The angular reflection dispersion (Figure 3 c) under realistic conditions (*Δ* = 1) shows a dimming in reflection and higher *FWHM* as calculated in Figure 3 b whilst the overall angular dependency remains unchanged compared to the ideal case scenario (*Δ* = 0).

Interestingly, the reflectance reduction due to disorder is much stronger under low light since the photonic crystal is more well-defined and hence more sensitive to disorder. Moreover, colour production has been identified as a possible functionality for this type of natural photonic structure. Therefore, a reduction in the total reflectance might result the iridoplast (and therefore the full begonia leaf) to be less conspicuous for pollinators which was proposed as one possible biological function of this type of nanostructures in the first documented descriptions of iridoplast (20,29). However, the recent and more detailed description of iridoplasts makes it worthwhile to study the effects caused by disorder on the absorption of the organelle rather than the colour production alone.

Figure 3 d shows that, with the expected uncertainty (*Δ* = 1), light absorption is enhanced (*γ* > 1) under low light conditions for *λ* ≳ 485 nm with a peak at *λ* ≈ 515 nm. Under high light, absorption is enhanced (*γ* > 1) for *λ* ≳ 565nm with a peak at *λ* ≈ 600nm. This represents a negligible redshift from the uniform iridoplast (*Δ* = 0). Compared to the perfectly ordered simulations, the disordered case, under both light conditions and due to a smaller overall reflectivity, shows higher absorption at short wavelengths. At longer wavelengths, under both light conditions, the absorption decreases. Altogether, disorder flattened the peaks and troughs along wavelength the absorption enhancement distribution. The overall absorption enhancement caused by the slow light effects on the absorption remains nearly unchanged with *γ*_*total*_ = 1.06 and *γ*_*total*_ = 0.99 for low and high light conditions.

Our results suggest that the conclusions reached with the perfectly ordered scenario apply as well to realistic disorder but with minor differences on the reflectance absolute values. That is, the light dependent dynamic changes of the photosynthetic tissue might explain the reversible changes in structural colour and might also provide a way to tune light absorption even when naturally occurring variability is taken into account. Our results point towards a given robustness to disorder of the photonic properties of iridoplast which could represent an advantage for light absorption under certain illumination conditions.

### c. Photonic properties of bisonoplasts

In order to obtain a full picture on the effects that reported morphological changes of chloroplast can have over photonic photosynthetic structures, it is worth using our model to study the bisonoplast as well. Similar to iridoplasts, bisonoplasts are a specialized type of chloroplast that has been recently reported to arrange thylakoid membranes in a regular manner to produce structural colour and, most likely, to manipulate light for photosynthesis. Their photonic properties were recently characterized for the first time (32). A schematic of their structure is represented in Figure 4 a and it behaves as a one-dimensional photonic crystal which reflects strongly in the blue, around 450 nm (21,22,32,33).

**Figure 4.**
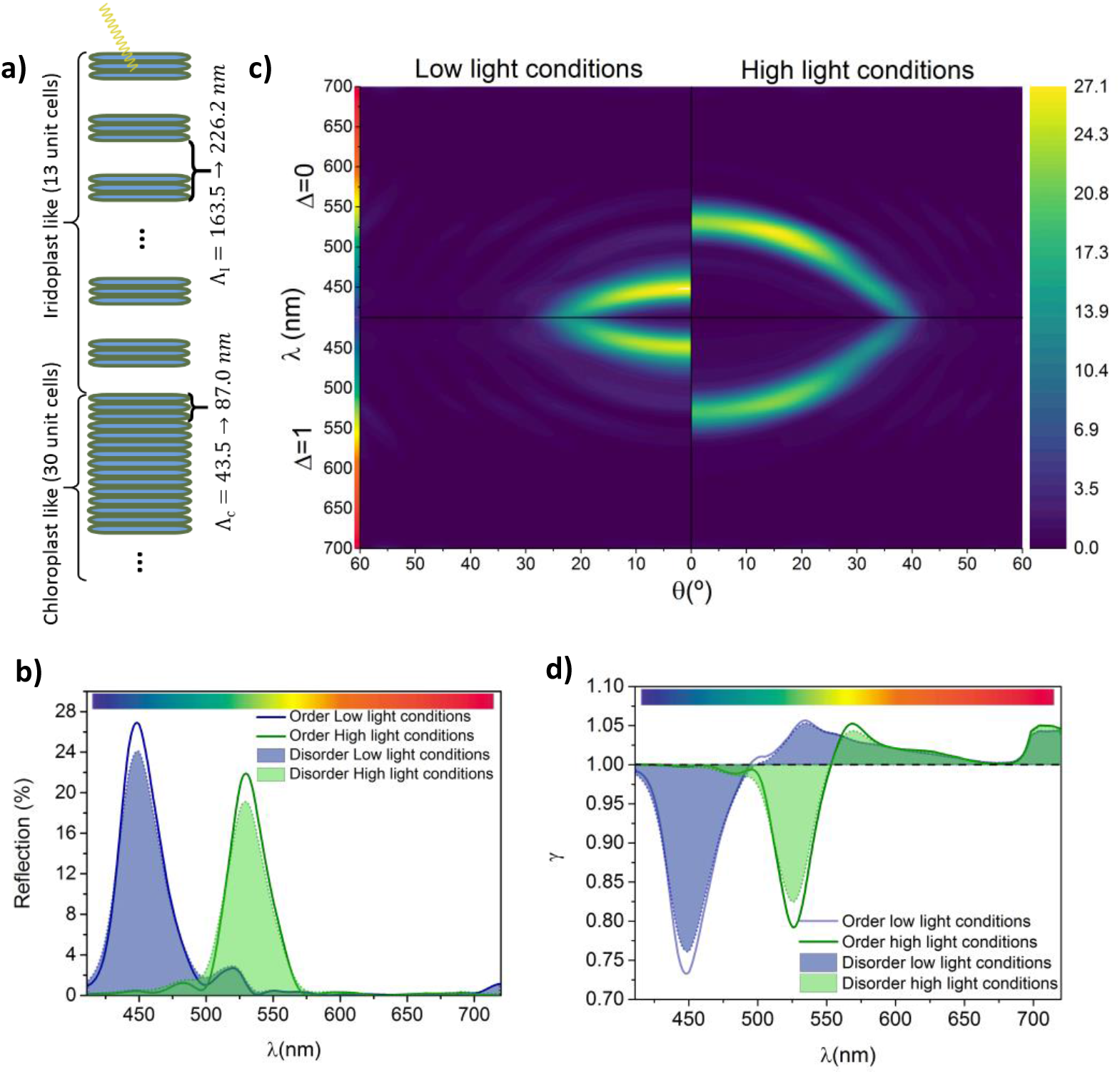
Photonic properties of bisonoplasts. **a)** Schematic of the structure of bisonoplasts. **b)** Normal incidence reflection spectrum of the bisonoplast at ordered (*Δ* = 0) and disordered conditions (*Δ* = 1) under low and high light conditions. **c)** Angular dependent reflection spectrum of the bisonoplast at *Δ* = 0 and *Δ* = 1 under low and high light conditions. **d)** Absorption enhancement (*γ*) of the bisonoplast under different order and light conditions.

In this model we consider 13 unit cells in the iridoplast-like part followed by 30 chloroplast like unit cells (33). Previously reported TEM measurements show no transition region between the iridoplast-like and chloroplast-like regions (33). We take the structural data measured from the bisonoplasts found in the plant species *Selaginella erythropus* (32). Even though the lattice length is similar to the iridoplast’s, the structure of the individual photosynthetic layers of these two photonic structures differs. We nonetheless assume that they react in a similar way to different light conditions. Table 2 summarizes the structural data used for these calculations. In our simulations, we consider ordered and disordered thicknesses in the same way as before. To calculate the absorption of the chloroplast in the absorption enhancement factor parameter of the bisonoplast *γ*(*λ*) = *A*(*λ*)_*bisonoplast*_/*A*(*λ*)_*chloroplast*_, we take the stromal gap (*d*_*s*_) between grana in the iridoplast-like phase to be zero (*d*_*s*_ = 0) leaving the rest of the structure unchanged.

**Table 2:** Average thickness (*μ*) and uncertainties (*σ*) of the different photosynthetic layers of bisonoplasts under different light conditions and the variation of each layer when going from dark to high light.

|  | Thickness under low light (nm) (36) | Variation going to high light (%) (12,25) | Thickness under high light (nm) |
| --- | --- | --- | --- |
| Thylakoid Membrane (M) | $4.5 \pm 0.6$ | $83 \pm 6$ | $3.7 \pm 0.6$ |
| Thylakoid Lumen (L) | $5.5 \pm 0.7$ | $196 \pm 40$ | $10.8 \pm 2.6$ |
| Stromal gap ($d_s$) | $120 \pm 5.7$ | $116 \pm 5$ | $139.2 \pm 8.9$ |

The model predicts that the reflection at normal incidence under low light conditions peaks around 450 nm (Figure 4 b), matching with previous experimental findings (32). Under high light conditions, the predicted structural changes would cause an 80 nm redshift in the reflection peak, having it at *λ* = 530 nm. There is a slight decrease in the absolute reflectance. To our knowledge, there hasn’t been to date any research on the reflection of the bisonoplast under different light environments and hence we can’t verify if this model matches the experimental reality. The angle dependent reflection (Figure 4 c) shows conclusions similar to the iridoplast over all angles: same central reflectance wavelength but slightly lower reflectivity and larger *FWHM* (Figure A 3). Just as for iridoplasts, the reflectance dispersion shows that under high light conditions we obtain a maxima around 25° rather than normal incidence. Again this is a direct consequence of the largest *n_t_* for *λ* < 500 nm.

This photonic structure was suggested to enhance light absorption in a similar fashion to the iridoplast (32). Under low light, we have an increase in the absorption (*γ* > 1) for *λ* > 505 nm and a peak absorption enhancement of only *γ* = 1.05 at *λ* = 535 nm. Under high light we have *γ* > 1 for *λ* > 550 nm and a maximum absorption enhancement of *γ* = 1.04 at 570 nm. Overall, the total absorption enhancement (*γ*_*total*_) in disordered conditions is of 1.01 under low light and 0.98 under high light, a much smaller effect than for iridoplasts. These findings directly contradict the hypothesis mentioned before (32). This is likely a consequence of the large number of layers in the bisonoplast which absorb most of the incoming light regardless of the slow light effect. Therefore, this effect has a much smaller contribution. Moreover, it is also possible that bisonoplasts might work on enhancing light absorption differently to iridoplasts, as it was suggested in ref. (33). From these results, we can also infer similar conjecture to the iridoplasts’ regarding the disorder: although they flatten reflection and absorption enhancement, the main conclusions are not affected by disorder.

### d. Enhanced absorption and biological importance

So far, we have analysed how differences in light conditions might affect the absorption of iridoplasts and bisonoplasts compared to chloroplasts. But it is also relevant to compare the changes in the absorption of the organelle under the different light conditions. For this, we define the single organelle dimensionless absorption enhancement factor as:

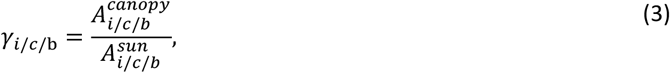

where 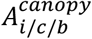 and 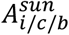 are the absorption of iridoplast (i), chloroplast (c) or bisonoplast (b) under low (canopy) and high (sun) light conditions respectively. Figure 5 shows *γ*_*i/c/b*_ using the morphological and refractive index parameters in Table 1 and 2. We use realistic data by including disorder to the thicknesses averaging over *S* = 50 simulations. To look further into the possible biological role of these dynamic structural changes, we calculate the total absorption enhancement parameter 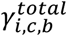 defined using the new *γ*_*i*_, *γ*_*c*_ or *γ*_*b*_ in Equation 1. We do this integration for both PAR at low (canopy) and high (sun) light conditions.

**Figure 5.**
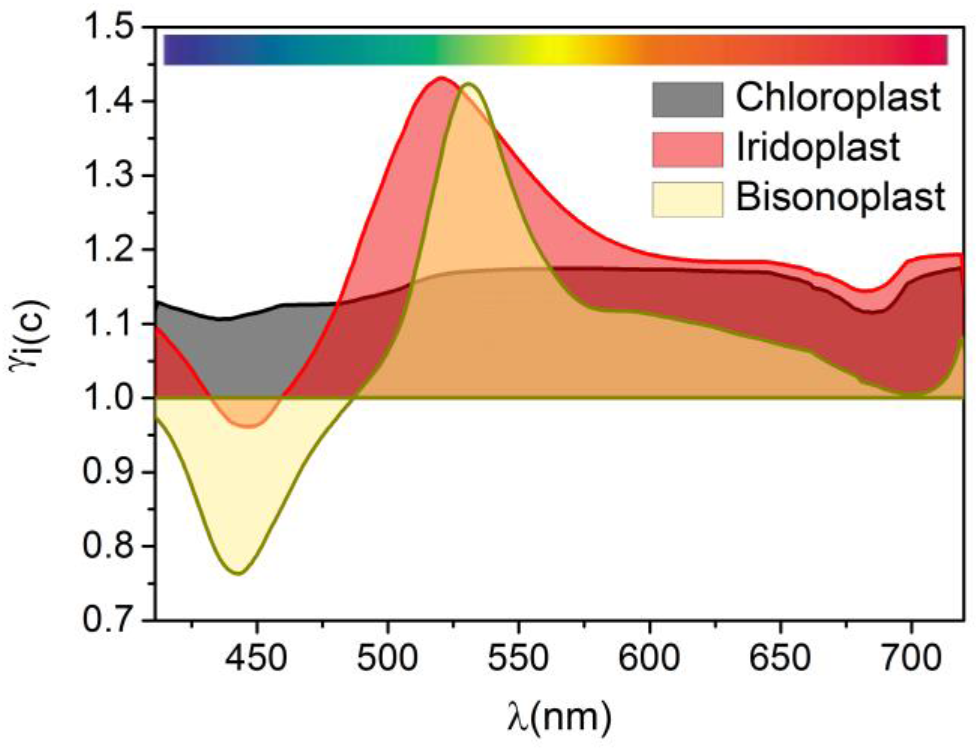
Light sensitive absorption tuning. Absorption enhancement (*γ*_*i/c/b*_) of the iridoplast/chloroplast/bisonoplast adapted to low light compared the same organelle adapted to high light conditions. Degree of disorder considered of *Δ* = 1.

Iridoplasts have a stronger absorption under low light conditions than under high light, with a peak at *λ* ≃ 520 nm where *γ*_*i*_ = 1.43. This enhancement is highly dependent on wavelength and for *λ* < 520 nm it starts to decrease due to a strong reflection. For chloroplasts we find a similar result, albeit one that is less wavelength dependent. Compared to high light, low light adapted chloroplasts have an absorption enhancement *γ*_*c*_ between 1.12 and 1.17, depending on wavelength. Bisonoplasts have the most wavelength dependent distribution with *γ*_*b*_ > 1 only for *λ* > 490 nm and having a peak of *γ*_*b*_ = 1.42 at *λ* = 530 nm. Integrating these results over the PAR shows that iridoplasts have *γ*_*total*_ = 1.19 and 1.25 above (direct sunlight) and below the forest canopy respectively. For chloroplasts, we obtain *γ*_*total*_ = 1.15 and 1.16 and bisonoplasts *γ*_*total*_ = 1.06 and 1.14 above and under the forest canopy respectively.

These results rise interesting points for discussion on the biological functions of the stratification of photosynthetic organelles. First, we obtain that all three organelle types, both under the canopy and at direct sun, enhance absorption for low light conditions (*γ*_*total*_ > 1). These results contradict those from ref. (3), where an expanded grana absorbs more light. This contradiction likely originates from the fact that the authors considered the grana to be an uniform bulk material, therefore neglecting the subwavelength stratification at thylakoid level which, taking into account their high refractive index for some wavelengths (Figure 1 d), can end up being relevant. Our results suggest that the three type of multilayer structures independently of their photonic (iridoplast and bisonoplast) or non-photonic (chloroplast) character have strong dynamic adaptations to better control light absorption depending on availability of light.

A second interesting conclusion is that for all three organelle types the total absorption enhancement is larger below the canopy than under direct sunlight 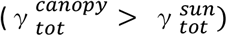. This means that a multilayer stacking and, in particular, a photonic crystal nanostructure represent a significant biological advantage specially developed for under the canopy environments where light is scarce but also spectrally different to the one available above canopy. In general, Figure 5 suggests that iridoplasts are the most efficient structure to enhance light absorption under the canopy with most of the spectral range in the *γ* > 1 region. Chloroplasts do not present a strong spectral dependence of the enhanced absorption under low light conditions as expected from their lack of photonic structuration. Yet, it should be noted that chloroplast present larger absorption under dark adaptation for the full PAR spectrum. Therefore, the documented grana contraction and expansion under different light conditions (Table 1) will allow chloroplasts to be more effective on capturing light under low light conditions according to our model. This is a relevant result expanding beyond the photonic photosynthetic organism. Oddly, bisonoplasts seem to present no advantage over chloroplasts since both 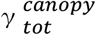 and 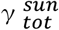 are smaller for the bisonoplast and hence the photonic structure seems to provide less light absorption control and no enhancement.

The experimental demonstration of our simulation results is extremely challenging but recent experimental data on the dynamics of iridoplasts and bisonoplasts could serve as a token for understanding other non-photonic photosynthetic organelles. The biological functionalities of the double stratification of the bisonoplasts remains unknown and numerical studies including 3D simulation might be needed for further understanding of light matter interactions within the organelle. The control of light absorption could be important to enhance photosynthesis when light is scarce and repress it when light is in excess, to avoid photodamage. Together with previous findings where expanded stromal lamellae induces photoprotection (17), this shows that chloroplasts, photonic and non-photonic, can manipulate the photosynthetic process to their advantage.

## 4. Conclusion

In this paper, we demonstrate that light dependent structural changes in the photosynthetic tissue of iridoplasts might explain the change in structural colour of *Begonia pavonina* leaves under different light conditions. It has been suggested that iridoplasts enhance light absorption at wavelengths longer than blue (λ > 480 nm). We showed that structural changes could also reduce this enhancement of light absorption. We have also looked into the effect of adding disorder to the system, by having layers with a varying thicknesses based on experimental data. This allowed us to consider a more realistic scenario which showed no significant differences from the perfect case. Similar simulations were performed on bisonoplasts which showed a reflectance red shift but a negligible absorption enhancement variation. The absorption enhancement of iridoplasts, chloroplasts and bisonoplasts under different light conditions shows that all systems have a very strong capability of tuning their light absorption, absorbing more under low light conditions. Iridoplasts have a much more sensitive change around green wavelengths, providing it a bigger absorption under the forest canopy, where they are found living, giving it a significant biological advantage. In these environments, iridoplasts allow for a better light absorption control than non-photonic chloroplasts. The model predicts a smaller light absorption enhancement for the bisonoplast compared to non-photonic chloroplasts meaning that the biological advantage of this photonic structure over a chloroplast remains unknown. Nonetheless, this work provides further evidence that it is important to consider nanophotonic effects when studying photosynthesis.

## Additional Information

Supplementary information and the data and Transfer-Matrix-Method simulation code for are available on request.

## Funding

This work was supported by the “Towards biomimetic photosynthetic photonics” project (POCI-01-0145-FEDER-031739) co-funded by FCT and COMPETE2020.

## Acknowledgment

We acknowledge Johannes Goessling for fruitful discussion.

## Disclosures

The authors declare no conflicts of interest.

## Author contribution

M.C. design and run optical models. M.C, W.P.W and M.L-G conceived the experiments and analysed the data. M. C., W.P.W and M. L. G. wrote the manuscript.

## Notes

### Competing Interest Statement

The authors have declared no competing interest.

