## Supplementary Material for "Light dependent morphological changes can tune light absorption in iridescent plant chloroplasts"

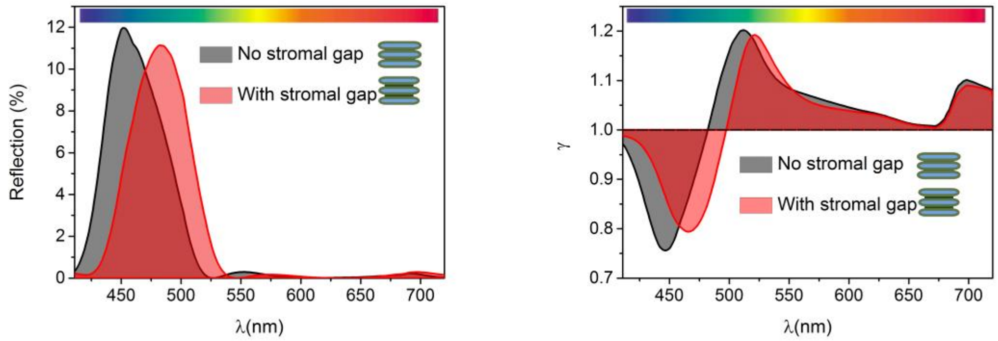


**Figure A 1. Simulations considering a stromal gap.** A 3.6 nm stromal gap between two consecutive thylakoid membranes was added for both light conditions and in the ordered situation. The rest of the structure remains unchanged. **a)** Normal incidence reflection spectrum. **b)** Absorption enhancement factor ($\gamma$) of iridoplast compared to chloroplast. Comparing the two photonic structure we find that there is a 30 nm redshift in reflectance. Slow light effects however show a much a less attenuated effect where the enhancement factor ($\gamma$) with stromal peaks at 520 nm, representing a 10 nm redshift compared to the simulations without this gap. These redshifts of the photonic properties are a consequence of the increase of the photonic lattice size. Considering the stromal gap, the total absorption enhancement factor $\gamma_{total}$ = 1.05, very similar when not considering the gap: $\gamma_{total}$ = 1.06. This thickness was taken from ref. (1) and it's likely an overestimate since in the iridoplast two thylakoid membranes and the stromal gap have a thickness of 6.7 nm (2) while in the former reference these amounted to 11.6 nm (4.0+4.0+3.6). With this, even considering an overestimation of the stromal gap, we obtain similar conclusions and it is therefore safe to consider the stromal gap negligible.

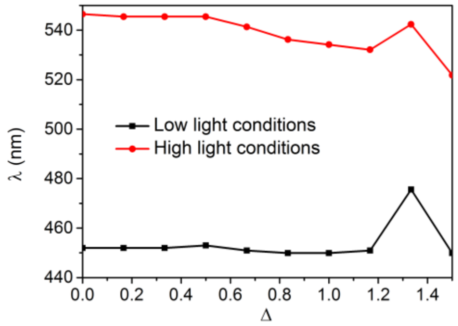


**Figure A 2. Effects of disorder on iridoplasts.** Reflection peak wavelength of the iridoplast at normal incidence for different disorder levels (i.e. varying $\Delta$). By inspection, we have a nearly constant reflection wavelength peak at low light along different levels of disorder and a blue shift of ≃10 nm for high light.

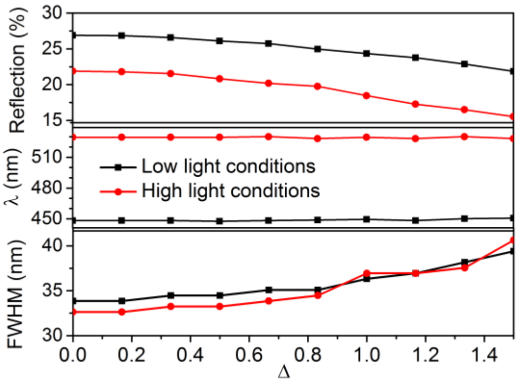


**Figure A 3. Effects of disorder on bisonoplasts.** From top to bottom: maximum absolute reflection, reflection peak wavelength and full width at half maximum (*FWHM*) at normal incidence for different disorder levels (i.e. varying $\Delta$).

Similar to the iridoplast, increasing level of disorder in the bisonoplast: reduces absolute reflectance, keeps the reflection peak wavelength unchanged and increases the *FWHM*.
